# Insights into rRNA processing and modification mapping in Archaea using Nanopore-based RNA sequencing

**DOI:** 10.1101/2021.06.14.448286

**Authors:** Felix Grünberger, Michael Jüttner, Robert Knüppel, Sébastien Ferreira-Cerca, Dina Grohmann

## Abstract

Similar to its bacterial and eukaryotic counterparts, ribosomal RNA maturation in archaea is a multi-step process requiring well-defined endo- and exoribonuclease activities. However, the detailed rRNA processing pathway in archaea remained elusive. Here, we employed long-read direct cDNA and direct RNA Nanopore-based sequencing to study rRNA maturation in three archaeal model organisms, namely the Euryarchaea *Haloferax volcanii* and *Pyrococcus furiosus* and the Crenarchaeon *Sulfolobus acidocaldarius*. Compared to standard short-read protocols, nanopore sequencing facilitates simultaneous readout of 5’- and 3’-positions, which is required for the classification of rRNA processing intermediates. More specifically, we i) accurately detect and describe rRNA maturation stages by analysis of terminal read positions of cDNA reads and thereupon ii) explore the stage-dependent installation of the KsgA-mediated dimethylations in *Haloferax volcanii* using basecalling and signal characteristics of direct RNA reads. Due to the single-molecule sequencing capacity of nanopore sequencing, we could detect hitherto unknown intermediates with high confidence revealing details about the maturation of archaea-specific circular rRNA intermediates. Taken together, our study delineates common principles and unique features of rRNA processing in euryarchaeal and crenarchaeal representatives, thereby providing a comprehensive picture of rRNA maturation pathways in archaea.

## Introduction

Ribosomes are universally conserved molecular machineries ensuring the efficient and accurate translation of mRNAs into proteins. Accordingly, the assembly of ribosomes is a key regulated housekeeping task enabling the synthesis of functional ribosomal subunits. Ribosome biogenesis, the process by which ribosomal subunits are assembled and matured, is a highly complex process requiring the coordinated action of several ribosome biogenesis factors, the RNA modification machinery, and ribonuclease activities responsible for the processing of the ribosomal RNA precursors (pre-rRNAs).

In contrast to bacteria and eukaryotes, little is known about the rRNA processing pathway in archaea (1–5). Our current knowledge suggests that the primary polycistronic rRNA precursor contains two processing stems formed by the 5’ leader and 3’ trailer sequences surrounding the 16S and 23S rRNAs (1, 4–7). In Euryarchaeota, the 16S and 23S rRNAs are additionally separated by the presence of an internal tRNA. In most archaea, the 16S and 23S rRNA processing stems contain a bulge-helix-bulge (bhb) motif, which is, in the context of intron-containing tRNAs, recognised by the splicing endonuclease endA (6–9). Similar to intron-containing tRNA maturation, processing at the bhb motifs is followed by the covalent ligation of the resulting extremities, thereby generating the archaeal-specific circular pre-16S and circular pre-23S rRNAs (1, 5, 7, 10, 11). The exact molecular mechanisms by which the circular pre-rRNA intermediates are further processed into linear mature rRNAs remain to be fully characterised (1, 5, 7).

Deciphering the rRNA processing pathway biochemically can be a time-consuming and tedious task (2, 12). Whereas nextgeneration sequencing has revolutionised sequencing-based approaches in RNA biology, understanding rRNA processing requires the analysis of 5’/3’ends co-occurrences, a feature that remains challenging to be determined by classical shortread sequencing technologies. Compared to standard shortread protocols, long-reads sequencing facilitates simultaneous readout of 5’- and 3’-positions, which is required for pre-rRNA stage classification of individual reads (13, 14). Additionally, single-molecule sequencing is a powerful tool enabling the detection of short-lived intermediates with high confidence. Accordingly, long-read sequencing strategies may facilitate obtaining rapid insights into pre-rRNA processing pathways in poorly studied organisms.

Here, we explored the combination of cDNA and native RNA sequencing to study archaeal rRNA maturation using the long-read sequencing technology offered by Oxford Nanopore. We tested the applicability of the ONT-based direct cDNA and RNA sequencing (i) to analyse pre-ribosomal RNA processing pathways and ii) to provide information on the timing of base modifications during rRNA maturation. We provide evidence that the long cDNA and RNA reads gathered on the ONT platform allow us to gain new insights into the poorly understood ribosomal RNA (rRNA) maturation pathway in various archaea. Moreover, we provide data that unravel the timely ordering of the KsgA-dependent dimethylations on pre-rRNAs in *H. volcanii*.

## Results

### Exploring rRNA processing and modification using long-read nanopore sequencing

To reveal characteristics of the rRNA processing pathway(s) in archaea we applied long-read nanopore sequencing to the total RNA of three archaeal model organisms, namely the Euryarchaea *Haloferax volcanii* and *Pyrococcus furiosus* and the Crenarchaeon *Sulfolobus acidocaldarius* (Figure 1A). For maturation stage classification, amplification-free direct cDNA sequencing was performed in a poly(A)-independent way using a custom 3’-adapter to improve the accuracy of 3’ end detection (Figure 1B,C,D), while direct RNA sequencing was used for stage-dependent base modification analysis (Figure 1B,E) (14, 21, 34). Note that we previously showed that artificial polyadenylation of prokaryotic RNAs impairs the accuracy of 3’ detection (14). Since RNA degradation influences read classification, samples for sequencing were selected based on the presence of rRNA peaks in the Bioanalyzer profiles after RNA pre-treatment and adapter ligation (Supplementary Figure 1A). During library preparation, RNAs were reverse transcribed using a custom primer. Successful reverse transcription was validated by size estimation of the cDNA product (Supplementary Figure 1B). Based on the length profiles, we conclude that some RNA species are prone to reverse transcription (RT) bias rendering absolute quantification impossible. This bias also applies to an *in vitro* transcribed 16S rRNA, which was used as a control and suggests that problems might occur particularly during RT of 16S rRNAs.

**Figure 1.**
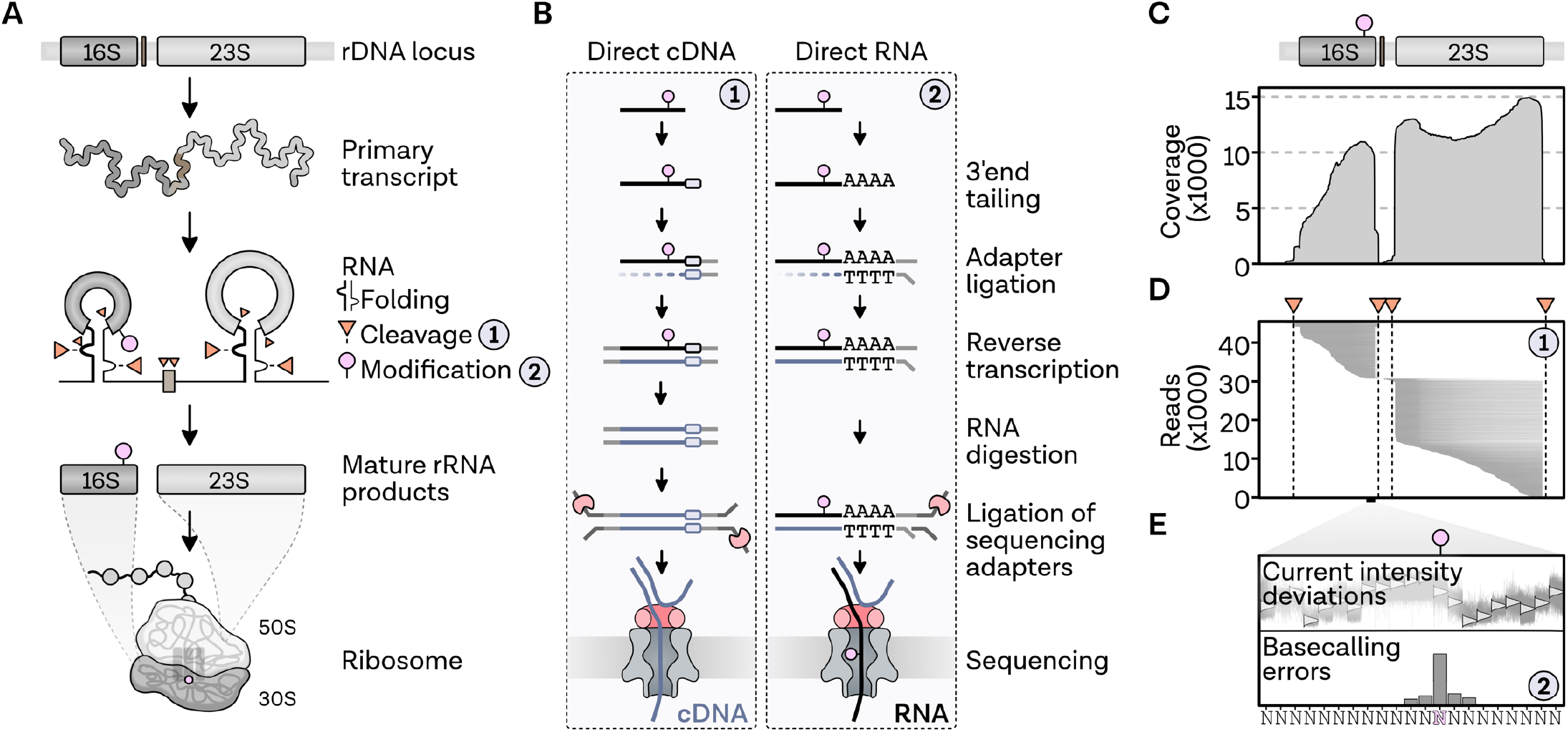
Exploring rRNA processing and modifications using Nanopore-based RNA-seq. **A,** Simplified pictogram of rRNA maturation in Archaea: The primary transcript containing the 16S (dark grey), internal tRNA (brown; only in Euryarchaeota) and 23S (light grey) is folded (folding indicated by bulge-helix-bulge structure), cleaved at multiple positions (orange triangles) and bases are modified (pink circle) until the mature rRNA products are formed that ultimately are an integral part of the ribosomal subunits (1, 5, 7). **B,** Nanopore-based library preparation methods for direct sequencing of cDNAs (left, 1) and RNAs (right, 2): For cDNA sequencing, 3’-adapter-ligated RNAs and custom VN-primers were used as input for the DCS109 protocol established by Oxford Nanopore. For RNA sequencing, libraries were prepared from artificially polyadenylated RNAs according to protocols RNA001 and RNA002 established by Oxford Nanopore. Adapters that carry the motor protein are highlighted with a red pacman symbol. **C,** Exemplary direct cDNA coverage profile and **D,** single-read plot of the rRNA locus 1 of *Haloferax volcanii*. Cleavage sites at the bulge-helix-bulge are indicated by orange triangles and dashed vertical lines. **E,** RNA base modifications can lead to deviations in the recorded current intensities from the expected signal and/or to systematic basecalling errors, which can be used to detect RNA modifications (34).

In total, about 90,000 to 270,000 reads were sequenced per total RNA library with a median q-score of 9.9 and a median read length of 1.2 kb (Supplementary Figure 2A-C). The length distribution of the sequenced reads corresponds to the expected sizes and enables the analysis of processing events at both extremities of the rRNA intermediates (Supplementary Figure 2D). After initial quality control, filtered reads (94.7 %, q-score filter: 7) were mapped to the reference genomes. About 99.5 % of the reads were successfully aligned with a median read length of about 200 bases shorter than the adapter-containing raw sequence length and a median identity of 92.9 %. In total, 71.7 % of the reads mapped to ribosomal RNAs of the three archaeal species. Coverage profiles of the *H. volcanii* samples indicated that about twice the number of reads mapped to the 23S rRNA sequence compared to the 16S rRNA although stochiometric amounts are expected to be incorporated into the ribosome. This unequal ratio was also observed or even more prominent in the other organisms (Supplementary Figure 3A). After analysing the distribution of aligned reads and mapped identities, we conclude that many rRNA-originating reads are potentially full-length reads, which are not influenced in their quality by length (Supplementary Figure 3B,C).

To prevent misidentification of terminal positions by random alignment of adapter sequences, full-length sequenced reads containing SSP and VNP adapters in the correct orientation were strand-oriented, and adapter-trimmed using pychopper. In total, 49 % of quality filtered reads contained the adapters in the correct orientation and were selected for subsequent analysis. Comparing the read coverage of full-length to unfiltered reads, we conclude that the most prominent coverage spikes in the unfiltered dataset are not genuine read ends and most likely caused by strand-switching problems (Supplementary Figure 4). Importantly, trimming allows using the soft-clipping parameter extracted from the CIGAR string as a meaningful parameter during the detection of pre-rRNAs characterised by spliced extremities, which will be described in the following paragraph (28).

### Insights into rRNA maturation in H. volcanii

After stringent filtering and quality control, we aimed to confirm and expand our knowledge on the poorly characterised multi-step ribosomal maturation process in archaea by performing enrichment analysis of terminal positions. The following analysis is illustrated using *Haloferax* as an example before applying the strategy to *Pyrococcus* and the Crenarchaeon *Sulfolobus*.

During the analysis, we noticed that some of the observed mature rRNA terminal positions did not match the available annotations at the NCBI databank (https://www.ncbi.nlm.nih.gov/genome/) (Supplementary Figure 5A-C). However, selected examination of the putative mature rRNA extremities obtained by direct cDNA sequencing did match our independent experimental validations by primer extension analysis of the 5’ ends of the 16S and 23S rRNAs of *H. volcanii* (Supplementary Figure 5D,E). These results suggest that positions based on our sequencing results most likely represent the genuine mature rRNA extremities. Therefore, the corrected annotation was subsequently used throughout the analysis (Supplementary Figure 5).

To re-trace the multi-step rRNA maturation process, we performed a co-occurrence analysis of read start and read end positions. Ribosomal RNA maturation in *Haloferax* starts with the transcription of a primary transcript from three promoters, which are also visible in a two-dimensional plot of read start and end positions (Figure 2A,B) (35). In accordance with the accumulated coverage data (Supplementary Figure 4), the most significant population of reads mapping to the 23S rRNA represent full-length 23S rRNA reads. In contrast, most of the 16S reads end at the mature 3’ end but have random 5’ starts. Processing at the predicted bhb positions, also visible in the scatter plot (Figure 2B), was analysed in more detail and is detected with single-nucleotide accuracy (Figure 2C,D). However, some of the reads starting or ending at the bhb have long soft-clippings originating from bases that are part of the sequenced read but not the alignment. Since the adaptors are trimmed off during pychopper filtering, these reads may correspond to the known circular precursors with covalently connected 5’ and 3’ bhb cleavage sites (7, 11). This is also supported by the fact that slightly divergent bhb positions can be explained by sequence similarities with the circularised rRNA.

**Figure 2.**
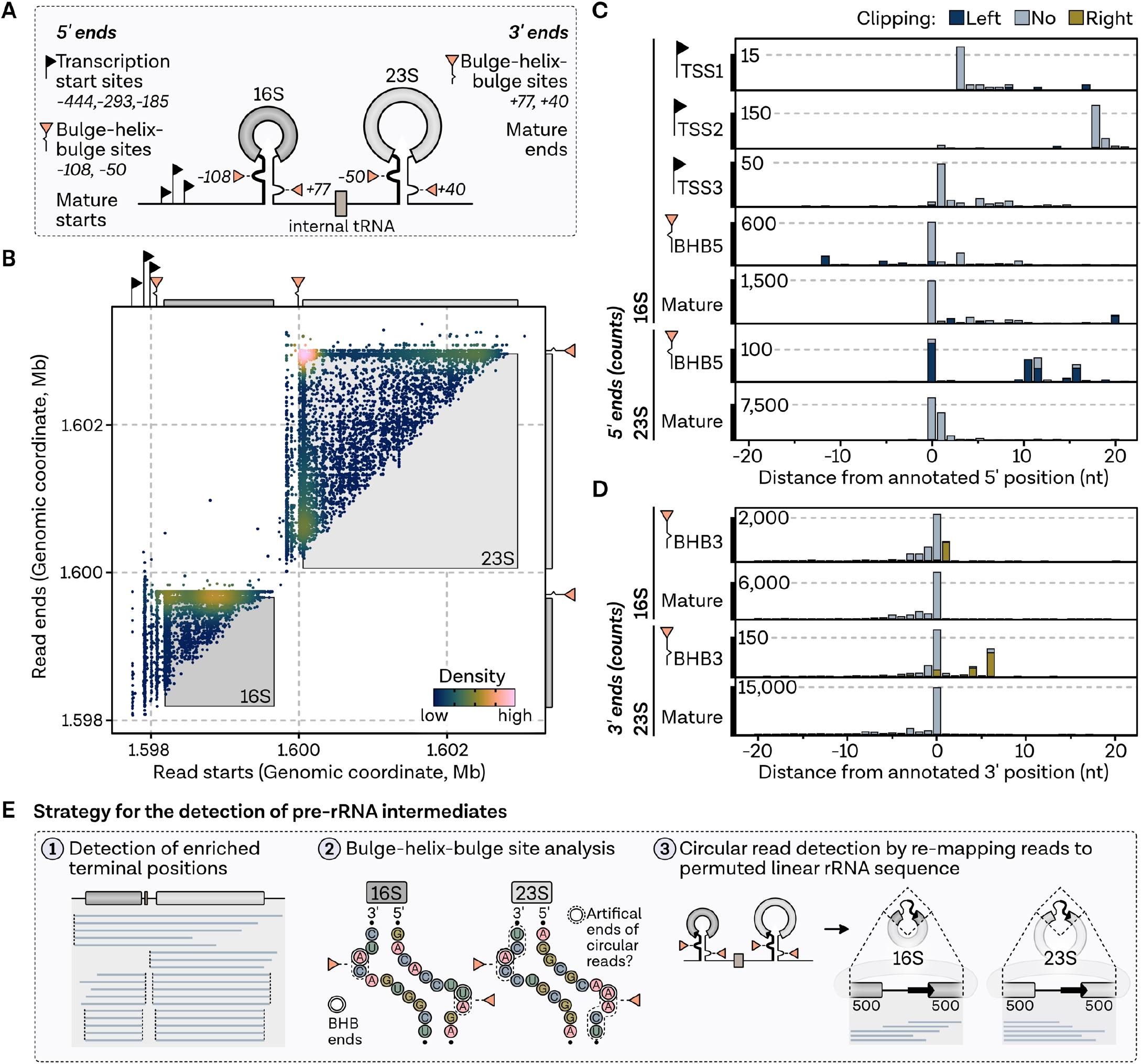
Strategy for the detection of pre-rRNA intermediates shown exemplarily for *Haloferax volcanii*. **A,** Expected 5’ and 3’ terminal positions of the 16S and 23S rRNAs based on previous publications (TSS: (35)), secondary structure prediction and previous analysis (bulge-helix-bulge: (11, 29)) and corrected positions of mature rRNAs (compare Supplementary Figure 5). **B,** Co-occurrence analysis of full-length cDNA read start (x-axis) and read end positions (y-axis) mapping to the rRNA locus 1 in *H. volcanii* (Hvo_3038-3041). Each read is shown as a single point, coloured by 2D density. Expected terminal positions are highlighted by black (transcription start sites) and orange (bulge-helix-bulge sites) triangles. **C,** Number of detected 5’ and **D,** 3’ ends with respect to the expected positions. Negative distance indicates upstream-mapping of 5’ or 3’ ends. Bars are coloured according to the soft-clipping profile in light blue (≤ 20 nt), dark blue (> 20 nt, left) and olive (> 20 nt, right). **E,** Strategy for the detection of rRNA intermediates: After the detection and co-occurrence analysis of enriched terminal positions (1), reads are analysed with respect to ends mapping to bulge-helix-bulge ends (2). Potential circular reads are detected by re-mapping reads to permuted linear rRNA sequences, designed by joining the bulge ends of the 16S and 23S rRNA.

To investigate this in more detail, Nanopore reads were remapped to a permuted RNA sequence designed by joining the 3’ bulge with the 5’ bulge of the 16S and 23S rRNA, respectively, to mimic the actual permutated sequence expected for circular pre-rRNAs (Figure 2E). For both, the mapping data based on the genomic sequence and the linear permuted sequence, pre-rRNA-related intermediates were selected based on read coordinates, abundance, biological interpretability, and prior characterisation, when available. Finally, the overall findings were used to extract a rRNA maturation pathway model summarised in Figure 3A (1, 5, 11).

**Figure 3.**
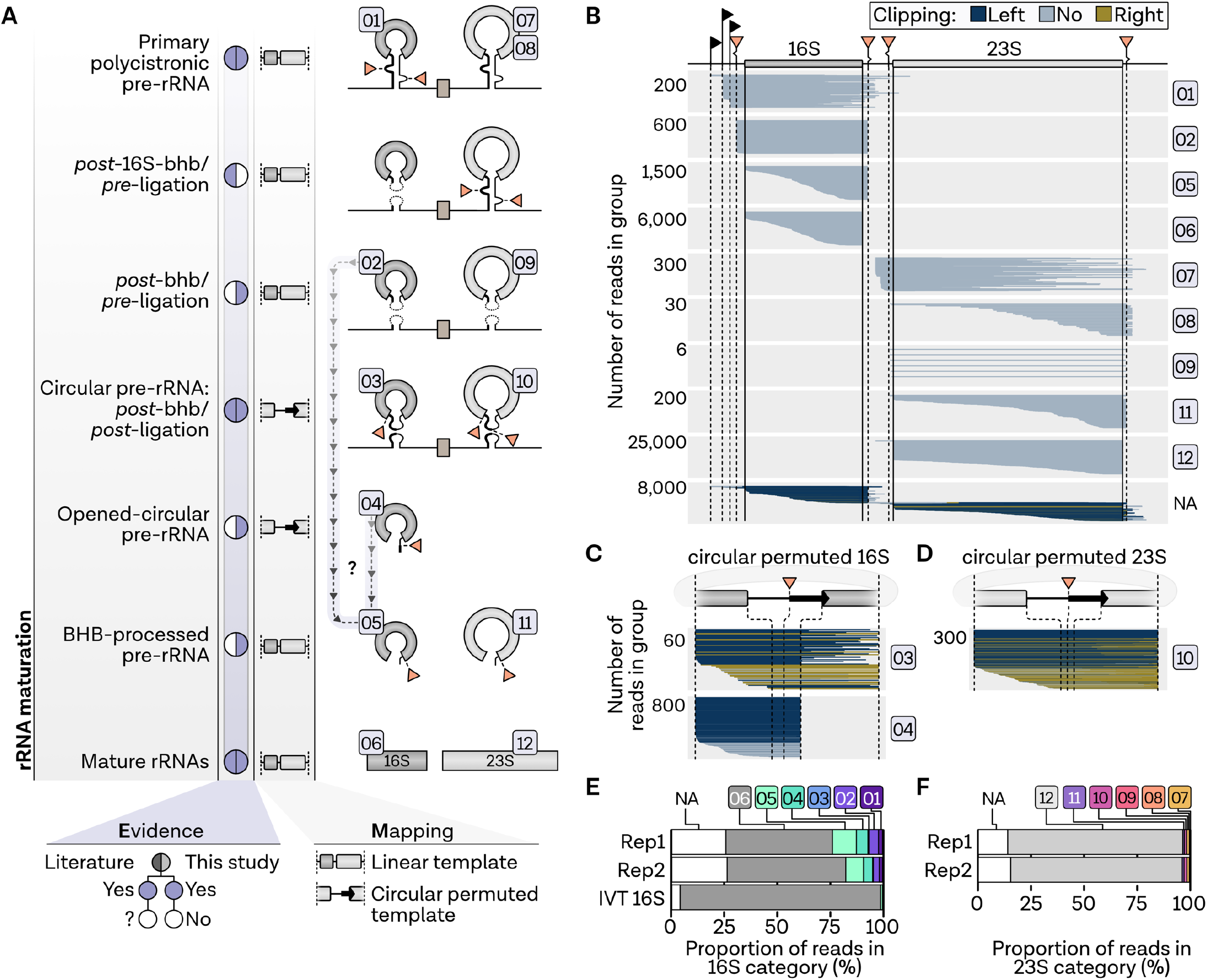
rRNA processing in *Haloferax volcanii*. **A,** Proposed rRNA maturation pathway in *Haloferax volcanii*.**B,** Single-track analysis of reads mapping to genomic linear template, **C,** the circular permuted 16S sequence and **D,** the circular permuted 23S sequence grouped by rRNA stage. Each read is shown as a single line sorted according to the read start position, while the relative number is shown on the y-axis. Lines are coloured according to the soft-clipping profile in light blue (≤ 20 nt), dark blue (> 20 nt, left) and olive (> 20 nt, right). **E,** The proportion of reads of the different rRNA processing states in the 16S groups and the **F,** 23S read groups are shown as stacked bar charts.

Despite the high sequencing depth, we did not detect a fulllength precursor consisting of the 16S leading-16S-tRNA-23S-23S trailing sequences in *H. volcanii*, suggesting that the primary rRNA transcript is rapidly processed. Accordingly, the primary pre-rRNA was detected as 3’ - (class 1 and 7) or 5’ - processed (class 8) reads (Figure 3B). Previously, functional *cis*acting element analysis suggested that the 16S rRNA bulgehelix-bulge processing occurs prior to internal tRNA and 23S rRNA maturation, leading to a precursor containing the 5’leader/3’trailer ligation of the processed 16S pre-rRNA flanking regions linked to the nascent 23S pre-rRNA intermediate. (11). However, we did not observe this particular precursor in the direct cDNA datasets of *H. volcanii*.Nevertheless, based on the end profiles, we identified four other classes of putative intermediates. In the first class (class 2,9) the pre-rRNA boundaries of these intermediates match the previously described bhb processing sites located within the 16S and 23S rRNA processing stems, respectively (11). These intermediates extend from the 5’ to 3’ bulge cleavage sites, however, these extremities are not covalently ligated and may correspond to post-bhb cleavage/pre-ligation pre-rRNA intermediates. An exemplary verification of the 5’ boundary of the post-bhb cleavage/pre-ligation pre-23S rRNA analysed by primer extension is provided in Supplementary Figure 5E. The second class (class 3,10) correspond to permuted reads covalently connecting the 5’ and 3’ bulge cleavage sites, and are likely observed as the result of random nicking of the circular pre-rRNA intermediates during sample preparations and are categorised as post-bhb/post-ligation pre-rRNA intermediates (Figure 3C,D). These intermediates were verified by re-mapping Nanopore reads to a permuted sequence designed by joining the 3’ and 5’ processing site of the bulge. Similarly, we detected a third main class (class 4), which corresponds to a putative pre-16S rRNA intermediate showing an immature 3’ end, which is extended by the typical permuted spacers sequence previously observed in the circular-pre-16S rRNA. This topology possibly results from linearisation of the circular pre-16S rRNA intermediate at the mature 16S rRNA 5’ end (opened-circular-pre-16S rRNA). This putative pre-rRNA intermediate is relatively abundant in *H. volcanii* (6 % of all reads mapping in the 16S rRNA region (Figure 5E), and strikingly shows a non-random 3’ end extremity, matching with the linearisation of circ-pre-rRNA at the mature 16S rRNA 5’ end (Figure 5C). Another previously undescribed intermediate, which is even more abundant in the 16S categories (11 %), and also present in the 23S rRNA maturation pathway (0.7 %), is characterised by randomly fragmented 5’ ends and a precise 3’ end at the bhb cleavage site. However, a temporal classification of this bhb-processed pre-rRNA intermediate is not possible based on the terminal positions. Both an alternative pathway without circularisation (after class 2,9) and processing after opening at the 5’ end (class 4) could account for the formation of this pre-rRNA intermediate (Figure 3B). Compared to the other stages, all reads classified as mature 16S or 23S rRNA products are expectedly the most abundant (Figure 3E,F). To confirm the presence of the class 2, 4 and 5 pre-rRNA intermediates in *H. volcanii* we performed Northern blot analysis after RNase-H dependent cleavage to deconvolute and discriminate between these pre-rRNA intermediates. As shown in Supplementary Figure 6, the presence of the opened-circular pre-rRNA and class 2 or 5 are readily observed in *H. volcanii*.

**Figure 4.**
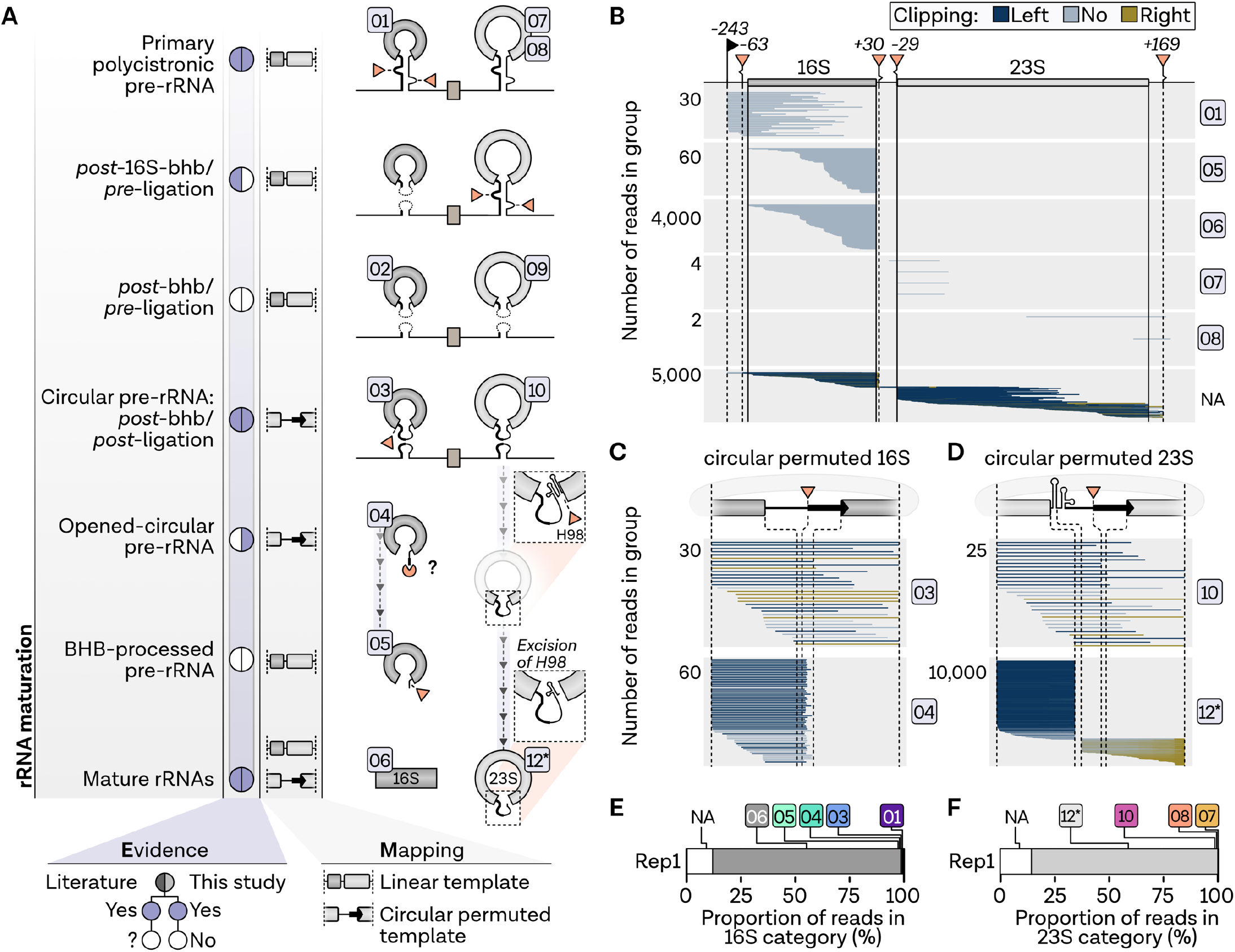
rRNA processing in *Pyrococcus furiosus*. **A,** Proposed rRNA maturation pathway in *Pyrococcus furiosus*. **B,** Single-track analysis of reads mapping to genomic linear template, **C,** the circular permuted 16S sequence and **D,** the circular permuted 23S sequence grouped by rRNA stage. Each read is shown as a single line sorted according to the read start position, while the relative number is shown on the y-axis. Lines are coloured according to the soft-clipping profile in light blue (≤ 20 nt), dark blue (> 20 nt, left) and olive (> 20 nt, right). **E,** The proportion of reads in the 16S groups and the **F,** 23S read groups are shown as stacked bar charts.

**Figure 5.**
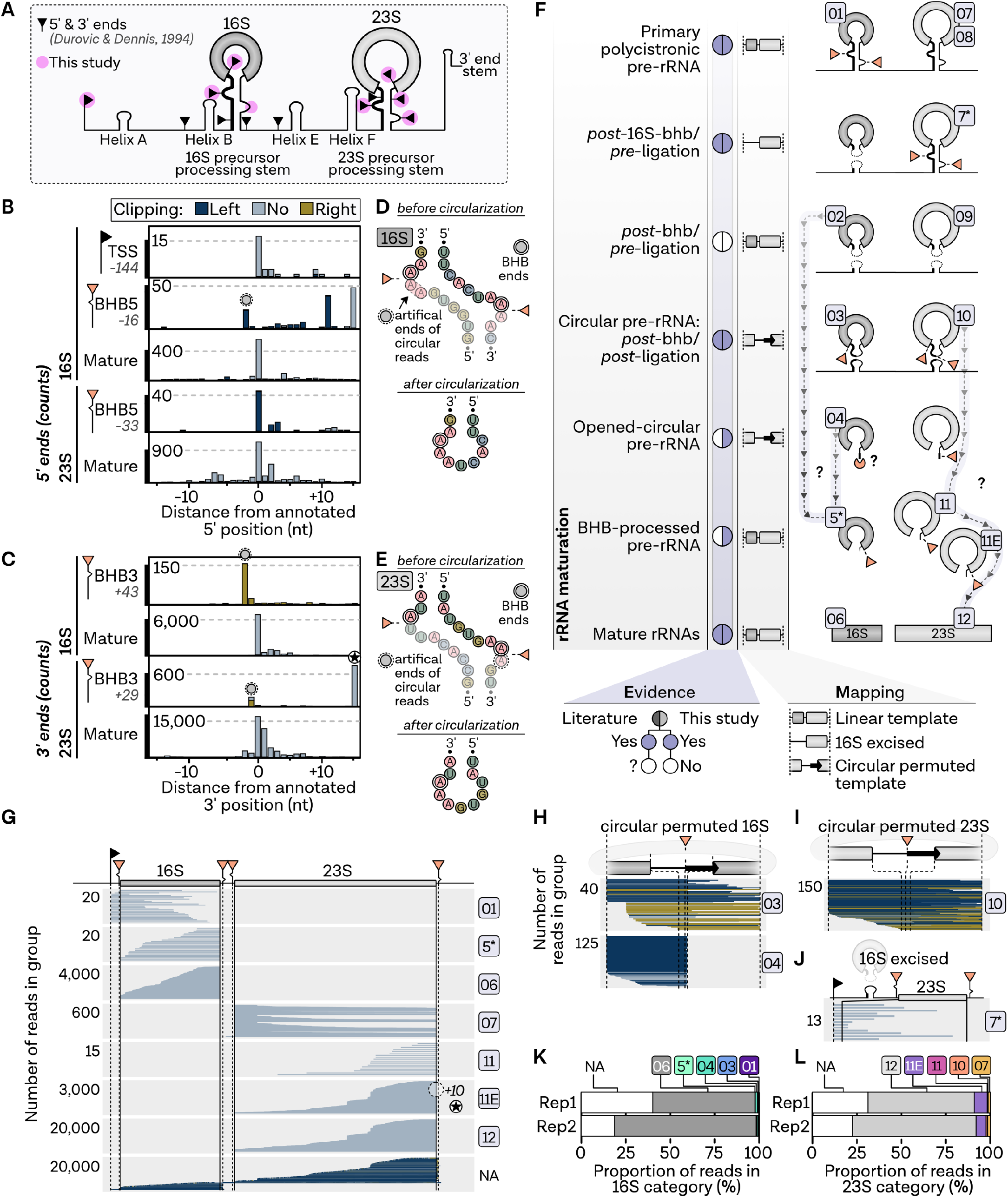
rRNA processing in *Sulfolobus acidocaldarius*. **A,**5’ and 3’ ends of mature rRNA in *Sulfolobus acidocaldarius* according to (38). Enriched ends found in this study are highlighted with pink circles. **B,** Number of detected 5’ and **C,**3’ ends with respect to the expected positions. Artificial ends of circular reads are highlighted by dashed circles. Negative distance indicates upstream-mapping of 5’ or 3’ ends. Bars are coloured according to the soft-clipping profile in light blue (≤ 20 nt), dark blue (> 20 nt, left) and olive (> 20 nt, right). **D,** Secondary structure of the bulge-helix-bulge sites of 16S and **E,**23S rRNA processing stems. Structures are shown before and after circularisation. **F,** Proposed rRNA maturation pathway in *Sulfolobus acidocaldarius*.**G,** Single-track analysis of reads mapping to genomic linear template, **H,** the circular permuted 16S sequence**, I,** the circular permuted 23S sequence and **J,** a linear template with excised mature 16S sequence grouped by rRNA stage. Each read is shown as a single line sorted according to the read start position, while the relative number is shown on the y-axis. Lines are coloured according to the soft-clipping profile in light blue (≤ 20 nt), dark blue (> 20 nt, left) and olive (> 20 nt, right). **K,** The proportion of reads in the 16S of the different rRNA processing states and the **L,**23S of the different rRNA processing states are shown as stacked bar charts.

### Insights into rRNA maturation in P. furiosus

Similarly, we determined the rRNA maturation stages in *P. furiosus* (Figure 4). In total, there are three deviations from the processing pathway observed in *Haloferax*:

First, opened-circular 16S pre-rRNAs (class 4) do not have a distinct 3’ end at the mature start position. This also applies to class 5, characterised by randomly degraded 3’ ends instead of a clear cut at the bhb-processing site. The third variation has been described recently and concerns the mature 23S rRNA (36). The special feature observed here is that the 23S rRNA is circularly permuted, which is a consequence of the excision of helix 98 after circularisation (Figure 4A). Thus, in this case, not only the pre-rRNAs (classes 3,10) but also the matured 23S rRNAs were detected by re-mapping to permuted sequences. Similar to the situation in *Haloferax*, the circular reads correspond to the covalent ligation of the 5’ and 3’ spacers generated by cleavage at the bulge-helix-bulge motifs within the processing stems (1, 9, 11).

To verify this hypothesis, we performed RNA structure prediction of the corresponding double-stranded RNA regions (Supplementary Fig. 7). In agreement with the permuted reads, we could place the corresponding extremities within the bulge-helix-bulge motifs. However, the 23S processing stem does not adopt a canonical bhb motif, but forms an alternative structure similar to the one previously described for the 16S rRNA bhb motif in *S. acidocaldarius* (7, 9, 37). Others and we could previously demonstrate that this alternative structure is compatible with circular-pre-16S rRNA formation in *S. acidocaldarius* (10, 11). Therefore, these permuted reads likely originate from random opening of the archaeal specific circular-pre-rRNA intermediates during sample preparation/sequencing (as observed for *H. volcanii*) and suggest that circular pre-rRNA intermediates are also produced in *P. furiosus* - a prevalent feature in rRNA processing found in many archaea. In fact, circularisation is a prerequisite for the subsequent maturation of 23S rRNA by excision of helix 98. Although our set-up does not allow absolute quantification, we find that the relative abundance of 16S and 23S rRNA precursors is lower compared to *Haloferax*.

### Insights into rRNA maturation in S. acidocaldarius

Next, *S. acidocaldarius* was analysed as a representative member of the Crenarchaeota. Regarding rRNA maturation, several features are homologous to the eukaryotic process. One of these features, termed site 0, is located about 70 nucleotides ahead of the 16S mature 5’ end. This feature was suggested to have similarities to the site A0 present in eukaryotic pre-rRNAs (Figure 5A) (38). Analysing the terminal positions in *S. acidocaldarius*, we did not detect site 0 to be enriched in our data sets (Figure 5B). However, this could be because it is most likely one of the earliest cleavage events resulting in a short-lived intermediate that is consequently underrepresented in the samples. Notably, we observed putative deviations in some of the bhb processing sites (Figure 5B,C). However, according to secondary structure analysis, this can be explained by sequence similarities after circularisation (Figure 5D,E).

We furthermore considered terminal positions described in (38) for our rRNA stage classification (Figure 5F). In addition to the truncated variants of primary RNA (class 1,7,8), we were also able to detect another intermediate that was already depicted in previous pathways but has not been discussed before (Figure 5G,J). This intermediate is a result of 16S processing prior to 23S processing, which leads to a post-16S-bhb RNA chimera (class 7*) characterised by a sequence with joined processed 16S leading and trailing sequences linked to the pre-23S rRNA. This state was confirmed by re-mapping of reads to a 16S excised sequence and is in good agreement with a pre-rRNA intermediate recently described based on *cis*acting element analysis in *H. volcanii* (11).

As observed for *Haloferax* and *Pyrococcus*, circularised forms of the 16S and 23S precursors could also be detected. However, we can find no circumstantial evidence to support the recent suggestion that the mature 23S rRNA is circular in *Sulfolobus* (36).

Regarding the other precursors, there are a few additional points worth mentioning: Similar to *Pyrococcus*, but different from *Haloferax*, the 3’ ends of the opened (class 4) and bhb-processed precursors (class 5) do not seem to be the product of a distinct endonucleolytic cleavage event. In contrast, we detected an additional cleavage event in the final steps of 23S rRNA maturation, which prominently accumulates in our data and was also described in other studies (class 11*) (Figure 5A,F,G,K) (38). In this case, processing occurs at position +10 in the 3’ region of the 23S rRNA, whereas the 5’ ends appear to be exonucleolytically processed as observed for the final rRNAs. In relative quantitative terms, this is also the only precursor found in substantial amounts (Figure 5K,L).

Taken together, our analysis confirms and expands the number of putative pre-rRNA intermediates in archaea. Moreover, this comprehensive framework provides an additional basis to facilitate further definition of common and specific principles of rRNA maturation in archaea.

### Towards stage-dependent analysis of rRNA modifications

In contrast to other sequencing techniques, Nanopore-based sequencing offers the possibility to detect base modifications directly as these modifications lead to an electric current signal that differs from the expected theoretical distribution obtained by the unmodified nucleotide sequence (30, 34, 39–42). In turn, these signal changes might also lead to differences in the basecalling profiles (e.g. systematic errors or a drop in basecalling quality) (34).

Accordingly, after determining rRNA processing stages via direct cDNA analysis, we aimed to explore the stage-dependent installation of the KsgA-dependent dimethylation of adenosine bases (m^6^_2_A) in *H. volcanii* using direct RNA sequencing (Figure 6). To this end, we analysed a wildtype sample and a deletion mutant of an archaeal KsgA/Dim1 homologue comparing basecalling, as well as signal features in the context of the m^6^_2_A modification at position A1450/A1451 (A1518/A1519 *E. coli* numbering)(20, 43, 44). Looking at the entire 16S region, helix 45, which contains the two predicted dimethylated bases, was the only region for which a substantial increase in basecalling errors was detected (Figure 6A-C). However, single-read analysis showed that the modifications in the wild type do not result in a uniform change in the profile suggesting a certain heterogeneity in this rRNA base modification population (Figure 6B). This is in line with a previous observation: primer extension data suggested that the KsgA/Dim1-dependent modification is heterogenous in *H. volcanii* (20). In addition to describing the percent Error of Specific Bases (ESB), which is used by the algorithm implemented in Eligos2 to detect base modifications based on the sum of alignment errors of direct RNA-seq data, two signal specific features were also compared, namely the mean dwell values and the mean signal time (Figure 6D,E) (30, 31). However, compared to the ESB analysis, no substantial differences were found between wild type and deletion strain in the m^6^_2_A sequence context. Hence, Eligos2 and ESB description were used to explore the potential to detect the introduction of the KsgA-dependent base modifications at different processing stages of rRNA maturation in *H. volcanii*. Therefore, reads were first sorted according to the main classes of mature and pre-rRNA intermediates described above for *H. volcanii*. Compared to the wildtype, bhb-processed pre-rRNAs (class 5) account for about twice as much of the total 16S reads in the KsgA-deletion mutant, suggesting that this late pre-rRNA class is accumulating in absence of KsgA (Supplementary Figure 8). Using Eligos2 we identified the stage-dependent installation of RNA modification with singlebase resolution (Figure 6G) (30, 31). It is striking that by far the largest value of the odds ratio, which represents the level of error of native sequence over the unmodified RNA, could be assigned to position A1450. In both the mature 16S (class 6) and the bhb-processed intermediate (class 5), the second highest value is accounted for the second site of methylation, position A1451. Although there are some odds ratio outliers for early precursors due to lower read counts, no statistically significant modification could be detected for classes 1 and 2. The earliest timepoint for the KsgA-dependent modifications is thus after circularisation, which was confirmed by the ESB analysis (Figure 6H). Although there is little difference in the profiles of classes 3, 4, 5, and 6, the slightly higher ESB at position 1451 and the resulting odds ratio value could be indicative of actual dimethylations in the circular precursor.

**Figure 6.**
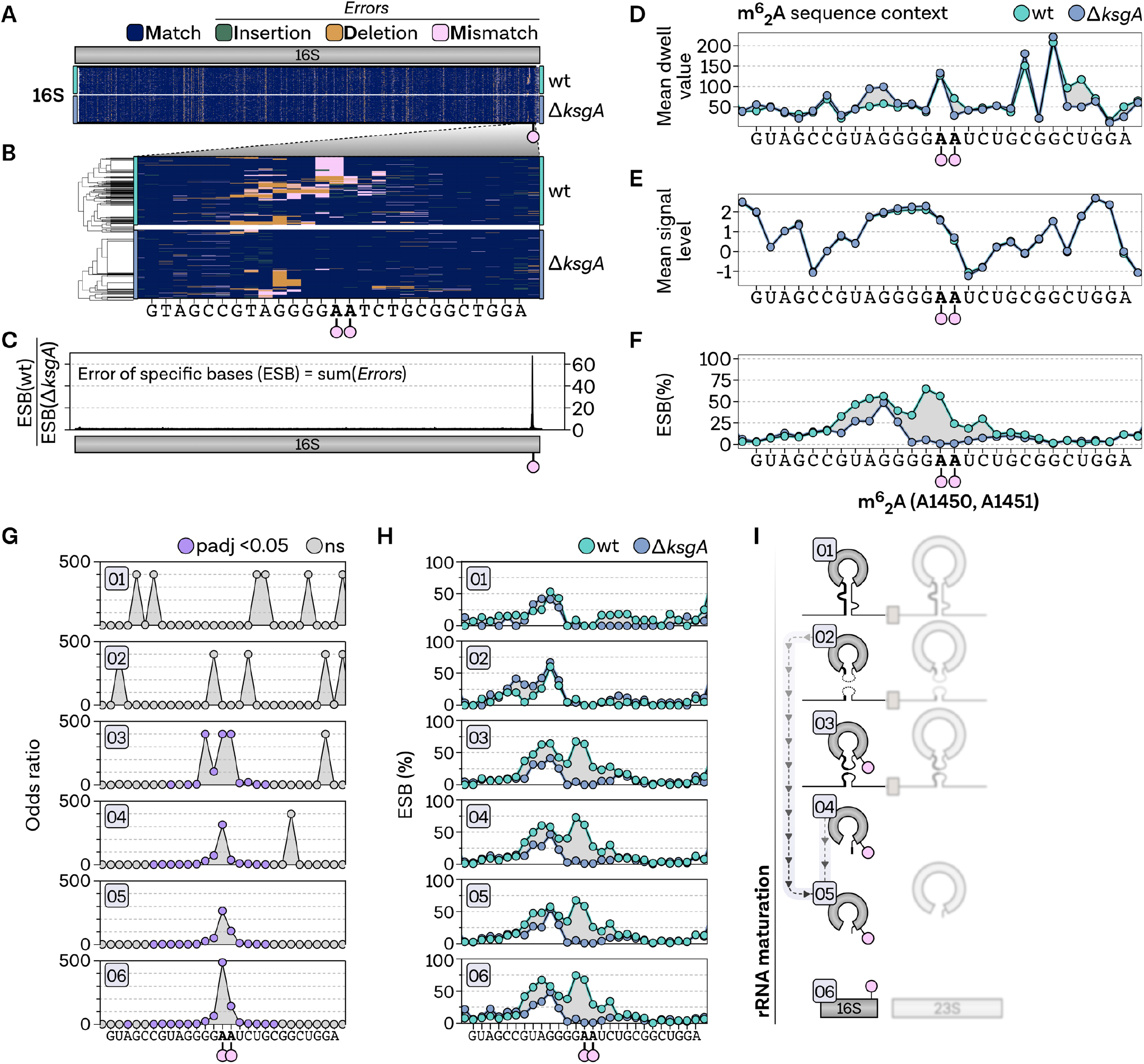
Direct RNA based modification analysis of KsgA-dependent dimethylations in *Haloferax volcanii*. **A,** Heatmap of single-read (y-axis) full-length wildtype (green) and Δ*ksgA* (purple) direct RNA reads coloured according to their mapping profile in blue (match), green (insertion), yellow (deletion) and pink (mismatch). **B,** Hierarchical clustering of reads mapping to helix 45 containing the KsgA-dependent dimethylation sites highlighted with pink circles. **C,** The error of specific bases (ESB) is calculated by the sum of all errors (insertion, deletion, mismatch) per position. The line plot shows the ESB of all reads from the wildtype sample divided by the ESB of the Δ*ksgA* sample. **D,** Tombo-calculated mean dwell time, **E,** signal time and **F,** the ESB are shown for the KsgA-dimethylations sequence context. Samples are coloured in green (wildtype) and purple (Δ*ksgA*), while the area between the lines is highlighted in grey. **G,** Odds ratios calculated by Eligos2 based on the comparison of wildtype and Δ*ksgA* reads grouped according to the categories used during the rRNA processing analysis. Significant positions are coloured in pink. **H,** ESB calculated for wildtype and Δ*ksgA* reads grouped according to the categories used during the rRNA processing state analysis. **I,** Proposed model for the stage-dependent installation of the KsgA-dependent dimethylations in *H. volcanii*.

Taken together, our analysis suggests that despite the current limitations, ONT allows to discriminate rRNA modifications across rRNA maturation events, which we included in our model of 16S rRNA maturation in *H. volcanii* (Figure 6I).

## Discussion

### Insights into rRNA processing in archaea

Ribosomal RNA processing proceeds via the coordinated and defined order of ribonucleases action (exonucleolytic and/or endonucleolytic cleavages), which generate pre-rRNA intermediates with defined premature rRNA sequences (2, 12, 45, 46). The establishment of faithful rRNA maturation maps in model organisms, like *E. coli, S. cerevisiae* or human cells has required numerous analyses over the past decades (2, 12, 45, 46), and remains a technical challenge. Therefore, methodologies that might accelerate the systematic analysis of rRNA maturation pathways across the tree of life need to be established to ultimately unravel the diversity of rRNA processing strategies in all domains of life. Beyond the identification of processing sites, the order of the processing events which can be, in part, deduced from co-occurrence analysis of the 5’ and 3’ extremities is of biological relevance.

In this study we could confirm and expand the presence of pre-rRNA intermediates and processing sites in the different archaeal organisms analysed, including the archaeal-specific circular-pre-rRNA intermediates (1, 5–7, 11). Together our findings are summarised into an updated archaeal rRNA processing model, which is discussed below:

The full length primary rRNA transcript was not identified in any of the archaeal organisms analysed. Similarly, this primary rRNA is generally difficult to observe in wildtype *E. coli* (47, 48). Collectively, these observations suggest that short-lived and/or low abundant pre-rRNA intermediates escape the detection capacity of the current experimental strategies. Accordingly, it is also difficult to infer differences in rRNA processing features between different (archaeal) organisms by virtue of observed pre-rRNA intermediates absence/presence pattern. In fact, these differences may also be related to organism-specific changes in pre-rRNA intermediates relative levels, which will depend on the sum of the reaction kinetics of the different maturation steps in a given condition.

Among the identified pre-rRNA intermediates, one precursor (class 7*), which is observed in *S. acidocaldarius* and includes ligation at the bhb motif of the upstream region of the 16S leader and downstream region of the 16S trailer sequences and continues to the downstream 23S sequences, is of particular interest. The presence of this ligation event suggests that the 16S rRNA bulge-helix-bulge processing occurs prior to 23S rRNA maturation. Although this ligation event was not identified by nanopore sequencing in *H. volcanii* and *P. furiosus*, this observation is in agreement with our recent functional *cis*-acting element analysis performed in *H. volcanii* (7, 11). In fact, based on this previous analysis, we have proposed a model by which 16S rRNA maturation proceeds and is required for the downstream maturation of the internal tRNA present in Euryarchaea and the 23S rRNA. Moreover, we have hypothesized that ligation of the 16S rRNA leader/trailer resulting from the 16S rRNA bulge-helix-bulge maturation process generates a putative new pre-rRNA intermediate for which the corresponding ligation event was observed using direct cDNA sequencing in an evolutionary distant archaeon (11).

The presence of circular pre-16S and pre-23S rRNAs and their processing sites could be verified and established in all organisms analysed in this study. Recently, we determined the functional requirement of the bulge-helix-bulge motifs for the formation of circ-pre-rRNAs in *H. volcanii* (11). Moreover, in analogy to intron containing-tRNA splicing, the rRNA bhb motifs are cleaved by the tRNA splicing endonuclease (endA) prior to covalent circularization (8, 9). Although intact circular RNA cannot be directly sequenced by the Nanopore approach, we noticed the presence of permuted transcripts by analysing clipping values, indicative of sequences that do not match to a linear template. Furthermore, most of the permuted reads of the circular rRNA classes 3 and 10 showed random and heterogenous 5’and 3’ends, thereby suggesting that these pre-rRNAs were likely the result of random nicking during sample preparation.

In addition to the circular pre-rRNAs, we observed pre-rRNA intermediates cleaved at the bhb motifs but not yet ligated to form circular pre-rRNAs in *H. volcanii (post*-bhb/*pre*-ligation pre-rRNAs; class 2,9). Whereas the presence of this intermediate processing step is theoretically expected, they were only detectable in larger numbers in the 16S maturation pathway of *H. volcanii*, suggesting that the maturation kinetics or stability of these pre-rRNA intermediates, prior to covalent ligation, varies among organisms and rRNA species.

Based on our current knowledge, several non-mutually exclusive hypothesis can be drawn for the final processing of circular pre-rRNA into linear mature rRNAs: (i) opening of the circular-pre-rRNA within the ligated spacer region and subsequent maturation of the 5’and 3’end; (ii) opening of the circular pre-rRNA by first maturation of the 5’end mature rRNA followed by 3’end maturation; or (iii) opening of the circular pre-rRNA by first maturation of the 3’end mature rRNA followed by 5’end maturation, as suggested earlier in a Methanogen (37).

Putative 16S pre-rRNA intermediates extended in its 3’ end by the presence of the ligated 5’ and 3’ spacers normally observed in the circular pre-16S rRNAs (class 4), may provide some indications how linearization of the circular pre-16S rRNA is achieved. This particular configuration is indicative of 5’end maturation of circular-pre-16S rRNA prior to final 3’end maturation, thereby generating opened-circular pre-16S intermediates. Additional properties of this putative intermediate are in agreement with its positioning during rRNA maturation and with the prevalence of 16S rRNA 5’maturation prior to its 3’end previously observed in bacteria and eukaryotes (2, 45). After 3’ processing, our data suggest that there may well be organism-specific differences that lead to the final product through the combination of endo- or 3’ -5’ exoribonucleolytic pathways. In particular, bhb-processed precursors (class 5) remain difficult to classify, as they could also represent 3’-extended early precursors. Noteworthy, these intermediates were also detected by our Northern blot analysis. Moreover, we could identify a new conserved ribonuclease required for the processing of class 4/5 pre-rRNA intermediates, suggesting that this step is conserved and might be characteristic of several archaeal rRNA processing pathways. The structure-function analysis of this ribonuclease will be reported elsewhere.

Overall, future functional characterisation of the *cis*- and *trans*-acting elements required for pre-rRNAs maturation will be necessary to further refine our general view on archaeal rRNA processing. In conclusion, despite some intrinsic limitations, we provide evidence that full-length rRNA sequencing can be a useful tool to approach intricated maturation pathway like rRNA maturation and expand our understanding of RNA maturation.

### Towards the mapping of stage-dependent installation of rRNA modifications

RNA modifications have been described already in the 50-60s, and have gained significant attention over the last years under the generic term of the epitranscriptome (49–51). However, the high-throughput analysis of these post-transcriptional modifications remains challenging and mostly relies on indirect methods, e.g. primer extension stops analysis and/or chemical recoding/derivation strategies (52, 53). Moreover, determining the timing of addition of rRNA modifications during rRNA processing remains challenging. Native RNA sequencing may fill an important gap to systematically analyse RNA modifications on a single-molecule level. However, global strategies enabling the faithful determination of RNA modification identity, position and timing of addition needs to be developed. Several recent analyses have explored different strategies to evaluate the capacity of nanopore sequencing to accurately detect RNA modifications (e.g. m^6^A) (13, 30, 34, 39–41, 54, 55).

We used a KsgA deletion strain as a background model, to analyse the installation of the almost universally conserved KsgA-dependent dimethylations (m^6^_2_A) at different stages of rRNA maturation in *H. volcanii* (41, 43, 44). By analysing basecalling profiles in wildtype and KsgA deletion strain we could unambiguously provide *in vivo* evidence that the archaeal KsgA-dependent methylations of the 16S rRNA are completed during the later stage of the small ribosomal subunit biogenesis in *H. volcanii* and may predominantly occur after circularisation of the pre-16S rRNA. These results are in good agreement with previous studies in eukaryotes and bacteria (20, 56–58).

Whereas nanopore sequencing may facilitate RNA modification analysis in general, the exact chemical nature of some modifications cannot be unveiled without prior knowledge and remain a challenging task which greatly benefits from the use of unmodified/hypo-modified reference sequences as suggested by a number of comparison-based algorithms (59–61). To facilitate high-throughput identification of RNA modifications, future studies will be required to develop and train algorithms improving the *de novo* identification confidence of diverse RNA/DNA modifications (62).

## Material & Methods

### Strains and growth conditions

*Pyrococcus furiosus* strain DSM 3638 cells were grown anaerobically at one bar excess of nitrogen in 40 ml SME medium (15) supplemented with 40 mM pyruvate, 0.1 % peptone and 0.1 % yeast extract at 95°C to mid-exponential phase and further harvested by centrifugation at 3,939 x g for 45 min at 4°C.

*Sulfolobus acidocaldarius* strain MW001 was grown in standard Brock medium (16–18) at 75°C to mid-exponential phase as described earlier (19).

Markerless deletion of *Haloferax volcanii* KsgA (Hvo_2746) was obtained earlier using the pop-in/pop-out procedure as described in (20). *Haloferax* strains (WT: H26) were grown in Hv-YPC medium at 42°C under agitation to mid-exponential phase as described previously (17, 20).

### *In vitro* transcription

First, genomic DNA was isolated from *H. volcanii* (H26) using standard phenol-chloroform extraction. The DNA was used as template for the PCR amplification of the 16S region using primers containing sites necessary for *in vitro* transcription using the T7 RNA polymerase (fwd: TAATACGACTCACTATAGGGATTCCGGTTGATCCTGCCGGA, rev: AGGAGGTGATCCAGCCGCAGA). The T7 RiboMax express kit (Promega) was used for the *in vitro* production of 16S rRNA under standard conditions.

### RNA isolation

For total RNA isolation, cell pellets were first lysed by the addition of 1 ml TRI Reagent (Zymogen). RNAs from *H. volcanii* and *S. acidocaldarius* were further purified using the RNA easy kit (Qiagen) according to manufacturer’s instructions. Alternatively, RNA from *P. furiosus* was isolated using the Direct-zol RNA Miniprep Plus protocol (Zymogen) following manufacturer’s instructions.

### RNA treatment for nanopore sequencing

To prevent secondary structure formation, RNAs were heat incubated at 70°C for 3 min and immediately put on ice before any of the following enzymatic treatments.

#### RNA treatment for direct cDNA sequencing

RNAs for direct cDNA sequencing were treated with T4 Polynucleotide Kinase (NEB, M0201) following the manufacturer’s non-radioactive phosphorylation protocol. After reaction clean-up using ZYMO RNA Clean & Concentrator-5 kit (ZYMO, R1013), 5’ triphosphates were removed enzymatically. To that end, 2 μg RNA were incubated for 30 minutes at 37°C with 2 μl RNA 5’ Pyrophosphohydrolase (NEB, M0356, 5,000 units/ml). The reactions were stopped by cleaning up the samples following the ZYMO RNA Clean & Concentrator-5 kit (ZYMO, R1013) protocol. To prevent artificial polyadenylation and improve 3’ end accuracy of nanopore sequencing, we used a custom 3’ cDNA RT primer. Accordingly, a custom 3’ adapter (5’-rAppCTGTAGGCACCATCAAT–NH2-3’, NEB, S1315S) was ligated to all RNAs, following the protocol described in (21). Briefly, 100 ng pretreated RNA was mixed with 50 pmol 3’ adapter, 2 μl 10× T4 RNA ligase reaction buffer (NEB, M0242), 10 μl 50% PEG 8000 (NEB, M0242), 1 μl 40 U μl^-1^ RNase Inhibitor (NEB, M0314), and 1 μl T4 RNA ligase 2 (truncated K227Q, NEB, M0242, 200,000 units/ml) and incubated at 16°C for 14 h. Finally, RNAs were cleaned up using the ZYMO RNA Clean & Concentrator kit (ZYMO, R1013).

#### Poly(A)-tailing for direct RNA sequencing

For direct RNA sequencing, RNAs were polyadenylated using the *E. coli* poly(A) polymerase (NEB, M0276) following a recently published protocol (22). Briefly, 5 μg RNA, 20 units poly(A) polymerase, 2 μl reaction buffer and 1 mM ATP were incubated for 15 min at 37°C in a total reaction volume of 50 μl. To stop the reaction and to remove the enzyme, the poly(A)-tailed RNA was purified with the RNeasy MinElute Cleanup Kit (Qiagen).

### Nanopore library preparation and sequencing

#### Direct cDNA sequencing

Libraries for direct cDNA sequencing were prepared following the instructions in the direct cDNA sequencing with native barcoding protocol (SQK-DCS109 with EXP-NBD104) from Oxford Nanopore Technologies with minor modifications. Briefly, the VN Primer was replaced with a custom 3’ cDNA RT primer (5’-ACTTGCCTGTCGCTCTATCTTCATTGATGGTGCCTACAG-3’, 2 μM). The size of the input RNA and the strand-switched cDNA was assessed via a Bioanalyzer (Agilent) run using the RNA 6000 Pico and the High Sensitivity DNA Kit (Agilent), respectively. During library preparation, the quantity and quality of the samples was tested using standard spectroscopic measurements (Nanodrop One) and using the Qubit 1X dsDNA HS assay and Qubit RNA HS assay kit (Thermo Fisher Scientific). Finally, samples were pooled in equimolar ratios, adapter-ligated, loaded onto a R9.4 flow cell (Oxford Nanopore Technologies) and sequenced on a MK1C device for 48 h.

#### Direct RNA sequencing

Libraries for nanopore sequencing were prepared from poly(A)-tailed RNAs according to the SQK-RNA001 Kit protocol (Oxford Nanopore, Version: DRS_9026_v1_revP_15Dec2016) with minor modifications. Briefly, Agencourt AMPure XP magnetic beads (Beckman Coulter) in combination with 1 μl of RiboGuard RNase Inhibitor (Lucigen) were used instead of the recommended Agencourt RNAclean XP beads to purify samples after enzymatic reactions. Also, the RTA adapter was replaced by custom adapters described in https://github.com/hyeshik/poreplex and reverse transcription (RT) was performed in individual tubes for each library. After RT reactions, cDNA was quantified using the Qubit DNA HS assay kit (Thermo Fisher Scientific) and equimolar amounts of DNA were used for ligation of the RNA Adapter (RMX) in a single tube. Subsequent reactions were performed according to the protocols recommended by Oxford Nanopore Technologies. Finally, libraries were loaded onto R9.4 flow cells (Oxford Nanopore Technologies) and sequenced on a MinION device for 48 h.

### Data analysis

#### Basecalling, demultiplexing and trimming of direct cDNA libraries

All fast5 reads from the direct cDNA sequencing run were basecalled using guppy (v. 6.3.2+bb5453e) in high-accuracy mode with a qscore cutoff of 7. Furthermore, basecalled files were demultiplexed in a separate step by guppy_barcoder using default parameters (barcode kit EXP-NBD104). For raw read quality control, relevant information was extracted from the MinKNOW sequencing summary file and the barcode summary file. Demultiplexed direct cDNA reads were strand-oriented and filtered for full-length sequenced reads containing the custom VN Primer and the strand-switching primer using pychopper with autotuned cutoffs and the recommended edlib backend (v.2.5.0). Additionally, protocol-specific read rescue was performed using the DCS109-specific option in pychopper. All full-length detected reads were merged and used for subsequent steps.

#### Basecalling, demultiplexing and trimming of direct RNA libraries

As some bioinformatic tools depend on single-read Nanopore files, we converted multi-read FAST5 files from the MinKNOW output to single-read FAST5 files using the ont_fast5_api from Oxford Nanopore (https://github.com/nanoporetech/ont_fast5_api). To prevent actual good-quality reads from being discarded (this issue was reported previously (23, 24)), we included both failed and passed read folders in the following steps of the analysis. Demultiplexing was done by poreplex (version 0.4, https://github.com/hyeshik/poreplex) with the arguments --trim-adapter, --symlink-fast5, --basecall and --barcoding, to trim off adapter sequences in output FASTQ files, basecall using albacore, create symbolic links to FAST5 files and sort the reads according to their barcodes. However, because of major improvements in the basecalling software, albacore files were not used. Instead demultiplexed FAST5 reads were basecalled using Guppy in high-accuracy mode (Version 6.1.3+cc1d765d3) without quality filtering. After basecalling and demultiplexing, artificially added polyA sequences were removed from the 3’ ends using cutadapt v4.1 (-a A{10}, -e 3, -j 0) (25).

#### Read alignment

Direct RNA and direct cDNA reads were mapped using minimap2 (v. 2.24-r1122) with standard parameters suggested for the alignment of noisy direct RNA reads (-ax splice -uf -k14) and Nanopore genomic reads (-max map-ont), respectively (26, 27). Reads were either aligned to the representative reference genomes downloaded from the NCBI or to circularly permuted sequences (described below).

In each case, --MD tag was used to include the MD tag for calculating mapping identities. Alignment files were converted to BAM files, sorted, and indexed using samtools (v.1.15.1) (28). Additionally, position-specific read coverages were calculated using samtools depth (-a -J). Read identities were calculated as previously reported (1-NM/aligned_read_length) after calculating the aligned read length by adding the number of M and I characters in the CIGAR string (14, 23, 28).

#### Detection of rRNA processing sites and classification of rRNA intermediates

Processing site detection was performed by enrichment analysis of start and end positions of reads mapping to the relevant rRNA regions. Sorting was achieved by classifying reads according to enriched and literature-expected terminal positions. For rRNA stage classification, coordinates of 5’ and 3’ ends, clipping information extracted from the CIGAR string, strand information and sequence identity were considered. Briefly, only reads with a mapping identity ≥ 90 % and with soft-clippings ≤ 20 nucleotides were used for classification based on linear templates (reads mapping to the representative genomes). Finally, stage-sorted reads were plotted in a genome browser-like view and evaluated based on the terminal positions. Enriched and previously undescribed combinations of connected 5’ and 3’ ends were considered in the classification.

#### Circular RNA detection

Circular reads were initially observed in a subset of reads, which end near/at the 5’-cleavage site of the bulge-helix-bulge, but are extensively left-clipped, which happens during mapping if the nucleotides further upstream do not match the 5’-leading, but the 3’-trailing region of the rRNA. The same is true for extensively right-clipped 3’ ends. Accuracy of 5’- and 3’-cleavage site detection using Nanopore reads was further evaluated by secondary structure prediction of the potential bulge-helix-bulge regions using RNAfold (29). To investigate circular rRNA reads in more detail, permuted linear sequences were created. These sequences contained 500 nt upstream of the annotated rRNA end to the predicted 3’cleavage site of the bhb site and were joined with the 5’-clevage site of the bhb up to 500 nt downstream of the annotated rRNA start. Nanopore reads were re-mapped to the linear permuted sequences and again categorised by their 5’ ends and 3’ ends as circular (reads cover unconnected mature parts of the rRNA and extend over the bhb) or opened-circular pre-rRNAs (read extends over bhb, 3’break at mature rRNA start). Additionally, reads potentially categorised as post-16S-bhb (only present in *Sulfolobus*, processing step 7*) were detected by mapping to a permuted sequence created by excising the 16S containing the leading and trailing sequences up to the bhb. For final stage classification, circular mapping reads were prioritized over linear templates with each read id only being assigned to one stage.

#### Modified base detection

For stage-dependent modified base detection direct RNA reads were sorted according to the read groups defined using the direct cDNA reads with two minor modifications. The quality threshold was lowered from 90 % to 80 % and the soft-clipping cutoff was increased from 20 to 30 nucleotides to account for the lower sequencing quality of direct RNA sequencing using Nanopore technology (13, 14).

Reads IDs were extracted for each stage and direct RNA fastq files sorted using seqtk subseq (v. 1.3-r106). Sorted Fastq files were subsequently mapped and used for calculating the frequency of correct, deleted, inserted and wrong nucleotides at each genomic position using pysamstats (v. v1.1.2). Note that the sum of substitution, deletions and insertion frequencies was defined as the Error of Specific Bases (% ESB) (30, 31). Additionally, modified base detection was performed using Eligos2 in pair_diff_mod (--oddR 0 --esb 0 --pval 1 --adjPval 1) to identify RNA modifications in the wildtype sample compared to the Δ*ksgA* sample (control sample) (30, 31). For modified base detection based on signal data, Fast5 files were first sorted to pre-determined rRNA stages using fast5_subset (ont_fast5_api). Next, multi-Fast5 files were converted to single read files using multi_to_single_fast5 (ont_fast5_api). Finally reads were pre-processed, resquiggled and the raw signals used to extract signal level and dwell value information using tombo (Version 1.5.1, https://nanoporetech.github.io/tombo). The tombo pipeline was run for pre-sorted files as well as for unsorted files (32). Downstream analysis was performed using custom R scripts (33).

### Primer extension analysis

Determination of 5’ ends of mature 16S and 23S rRNAs from *H. volcanii* by primer extension was performed as described previously (17). In brief, reverse transcription was performed with the indicated fluorescently labeled primers (oHv396: 5’-DY682-CCCAATAGCAATGACCTCCG for *H. volcanii* 16S rRNA 5’end; oHv622: 5’-DY782-GCTCTCGAGCCGAGCTATCCACC for *H. volcanii* 23S rRNA 5’end) and SuperScript III reverse transcriptase using 1 μg of total RNA as template. The resulting cDNAs and reference dideoxy-chain termination sequencing ladder reactions were separated on a denaturing 14 % TBE-Urea (6 M)-PAGE. Fluorescence signals (700nm and 800nm) were acquired using a Li-COR Odyssey system.

### RNaseH cleavage assay

5 μg of *Haloferax volcanii* total RNA were incubated with 10 pmol of oHv203 (AGGAGGTGATCCAGCCGC) and oHv254 (GTACTTCCCAGGCGGCTCG) and 1.5μl 10 x RNaseH buffer for 5 minutes at 65°C followed by an oligo annealing step for 10 minutes at 37°C in a total volume of 14 μl. 2 μl of RNaseH (10 U, NEB, M0297) was added followed by an incubation of 20 minutes at 37°C. 15 μl of 2x formamide buffer (99.95 % Formamide, 0.05 % Bromphenol blue) were added and the reaction stopped at 65°C for 15 minutes. RNaseH-cleaved total RNAs were loaded on a denaturing 1.5 % agarose gel and further analysed by northern blotting as described previously (19) using the following fluorescently labeled probes: Probe1, targeting the 16S mature rRNA (oHv707: DY682-GAGCTGGTGAGATGTCCGGC); Probe 2, targeting the 16S ligation site (oHv762: DY782-CCGTTCGGATaaggtgtccg); Probe 3, targeting the 16S 3’ trailer (oHv325: DY782-cgtgtgagccaccccgtcgg).

## Data availability

Raw direct RNA sequencing data sets (gzipped FAST5 files) have been uploaded to the Sequence Read Archive (SRA) and are available under project accession number PRJNA632538 (WT run: SRR11991303, Δ*ksgA* run: SRR11991308).

Direct cDNA data are available at the European Nucleotide Archive (ENA, https://www.ebi.ac.uk/ena) under project accession number PRJEB57168 (63).

## Code availability

Documentation and code of all essential analysis steps (tools and custom Rscripts) are available from https://github.com/felixgrunberger/rRNA_maturation.

## Acknowledgements

We thank all the members of the Ferreira-Cerca lab and of the Grohmann lab, especially Prof. Dr. Winfried Hausner and Dr. Robert Reichelt for fruitful discussions. We thank Katharina Vogl for her technical assistance. We furthermore would like to thank Jörg Soppa for discussion in the early phase of this project.

## Supplementary Figures

**Supplementary Figure 1.**
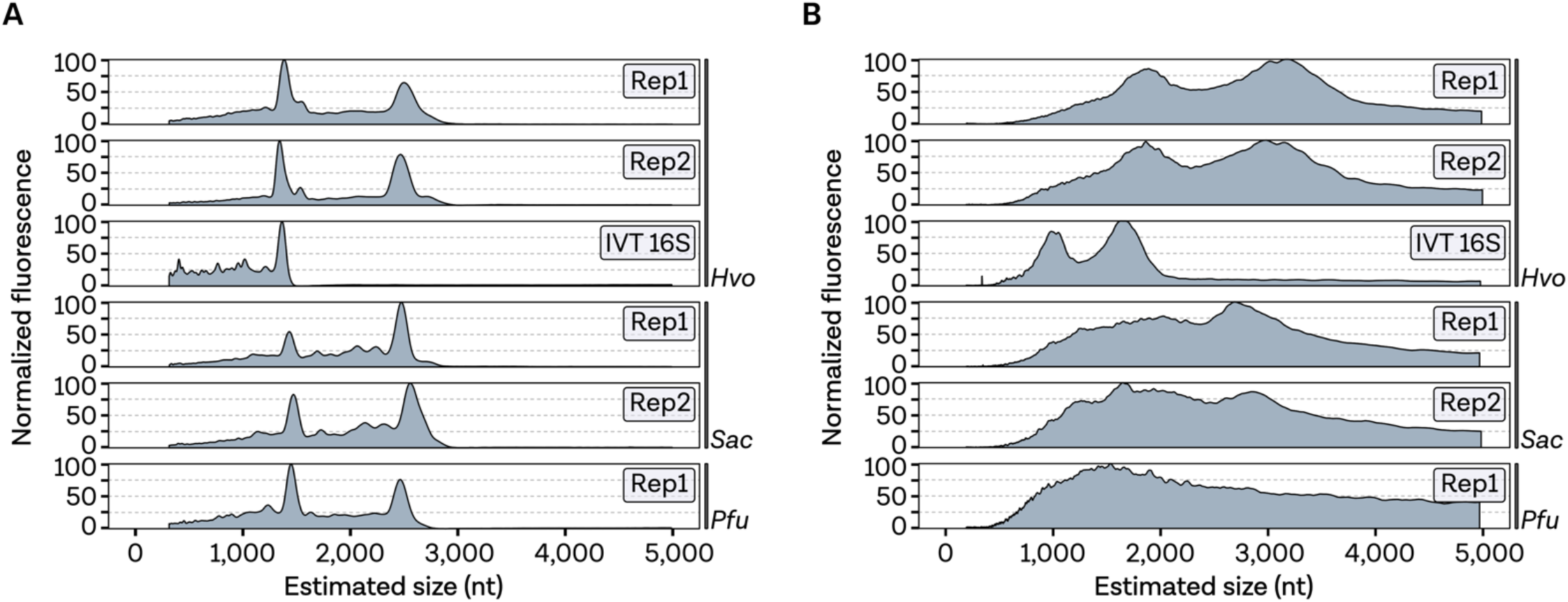
Bioanalyzer profiles. **A,** Bioanalyzer profile of pre-treated RNA used as input for the direct cDNA protocol and **B,** quality control of the cDNA during library preparation.

**Supplementary Figure 2.**
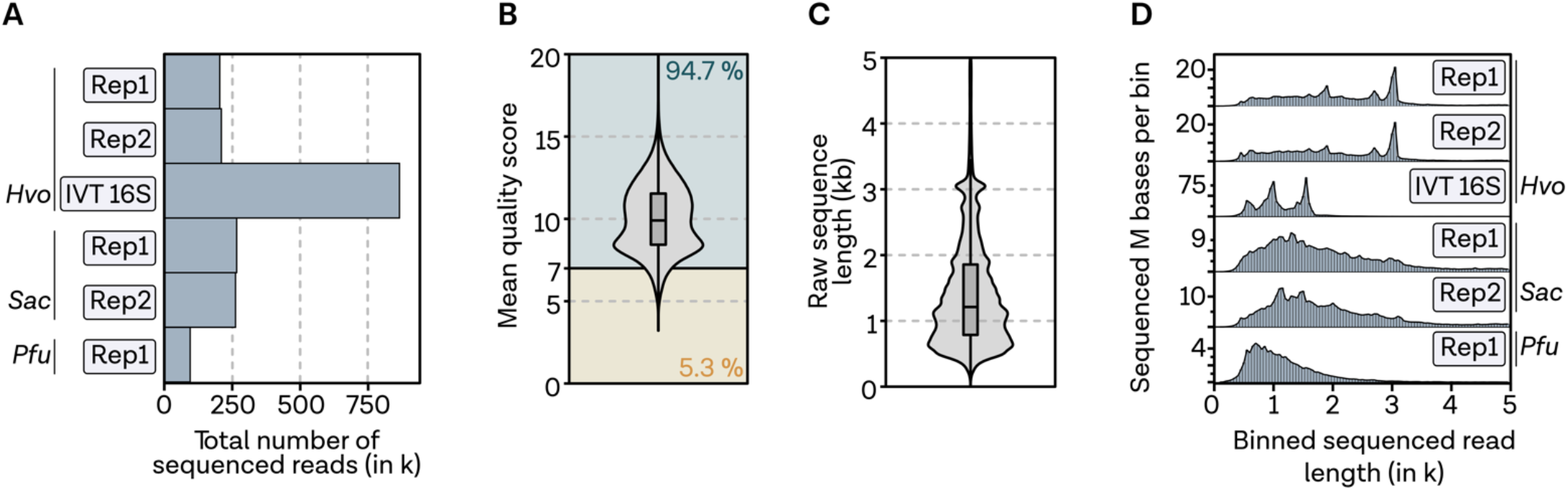
Quality control of sequenced cDNA reads. **A,** Total number of sequenced reads shown for each library. **B,** Mixed violin/boxplot showing the distribution of the quality scores (mean_qscore_template) and the number of reads passing and failing the quality filter of 7. **C,** Distribution of raw sequence length for all libraries containing total RNA. **D,** Distribution of binned sequence read length (bin size 50nt).

**Supplementary Figure 3.**
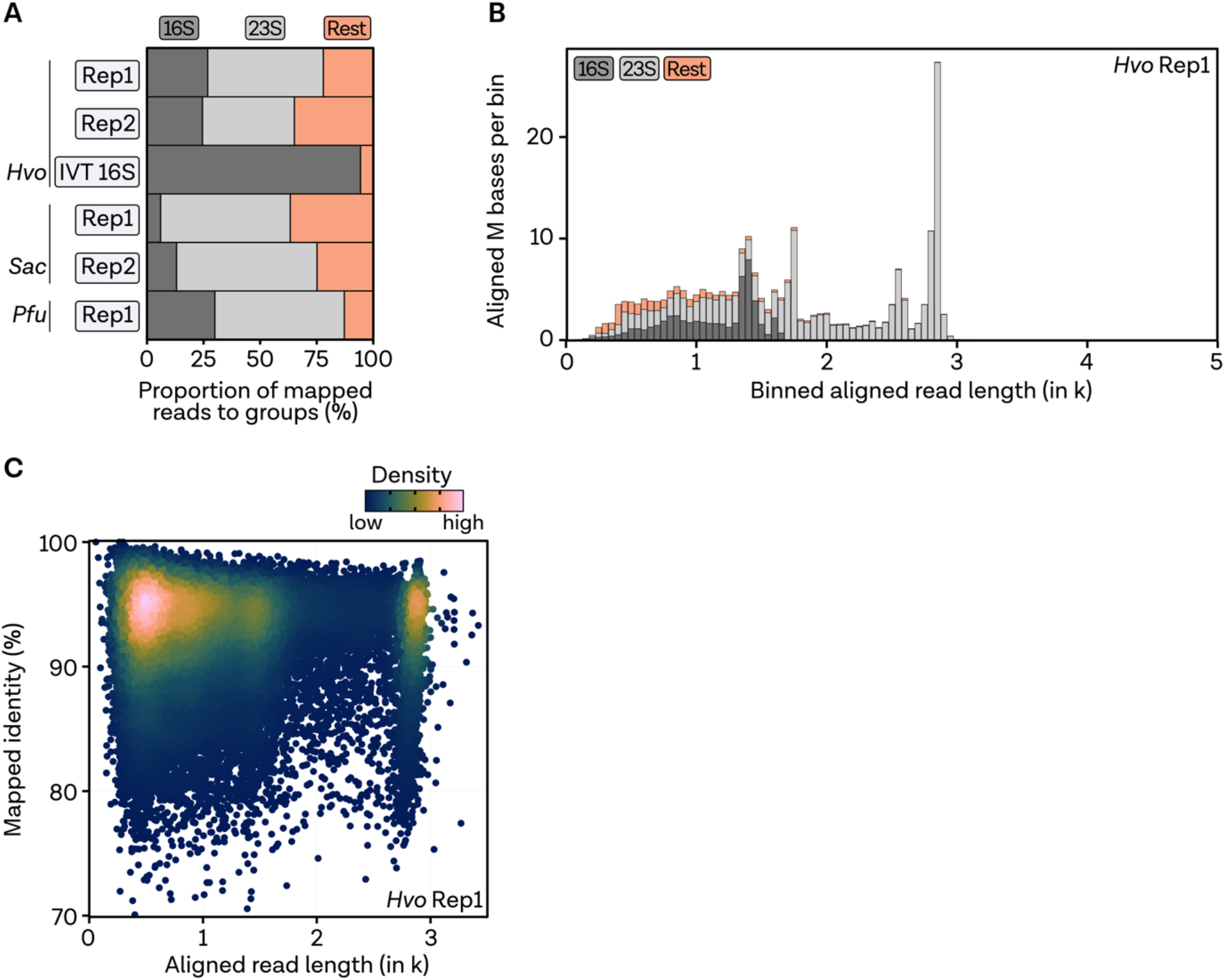
Quality control of mapped reads. **A,** Proportion of reads mapping to 16S (dark grey), 23S (light grey) and everything else (rest, white). **B,** Distribution of aligned read length (bin size 50nt), stacked by alignment group. Replicate 1 of *H. volcanii* is shown as a representative sample. **C,** Density plot of 20,000 randomly sampled reads showing the aligned read length to mapped read identity coloured by 2D density.

**Supplementary Figure 4.**
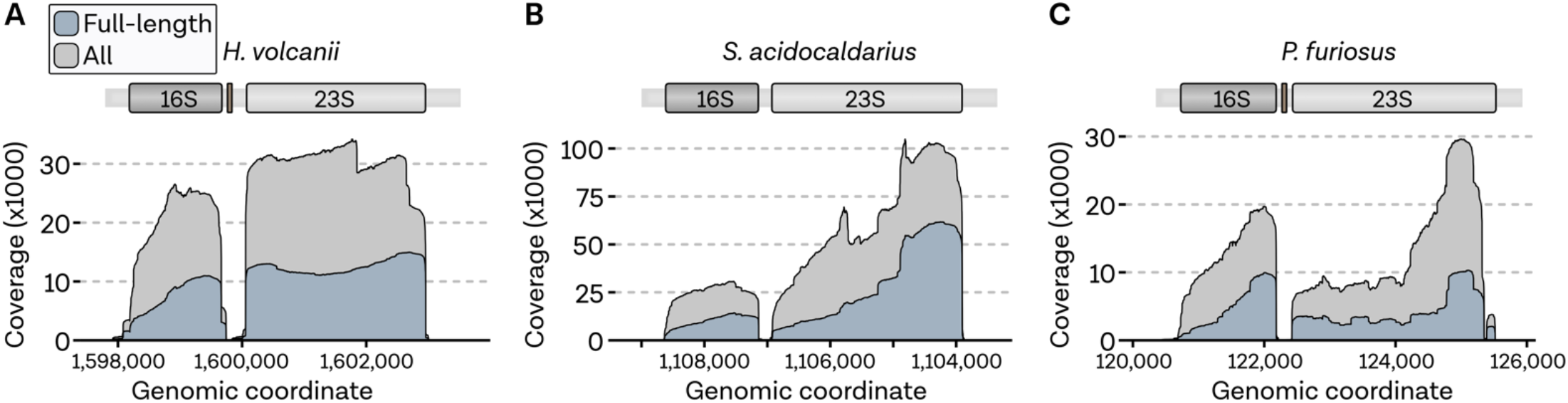
Coverage profile comparison of full-length and unfiltered reads. **A,** Coverage profiles of rRNA regions coloured by alignments from unfiltered (grey) or pychopper filtered full-length reads (blue) for *H. volcanii*, **B,***S. acidocaldarius* and **C,** *P. furiosus*.

**Supplementary Figure 5.**
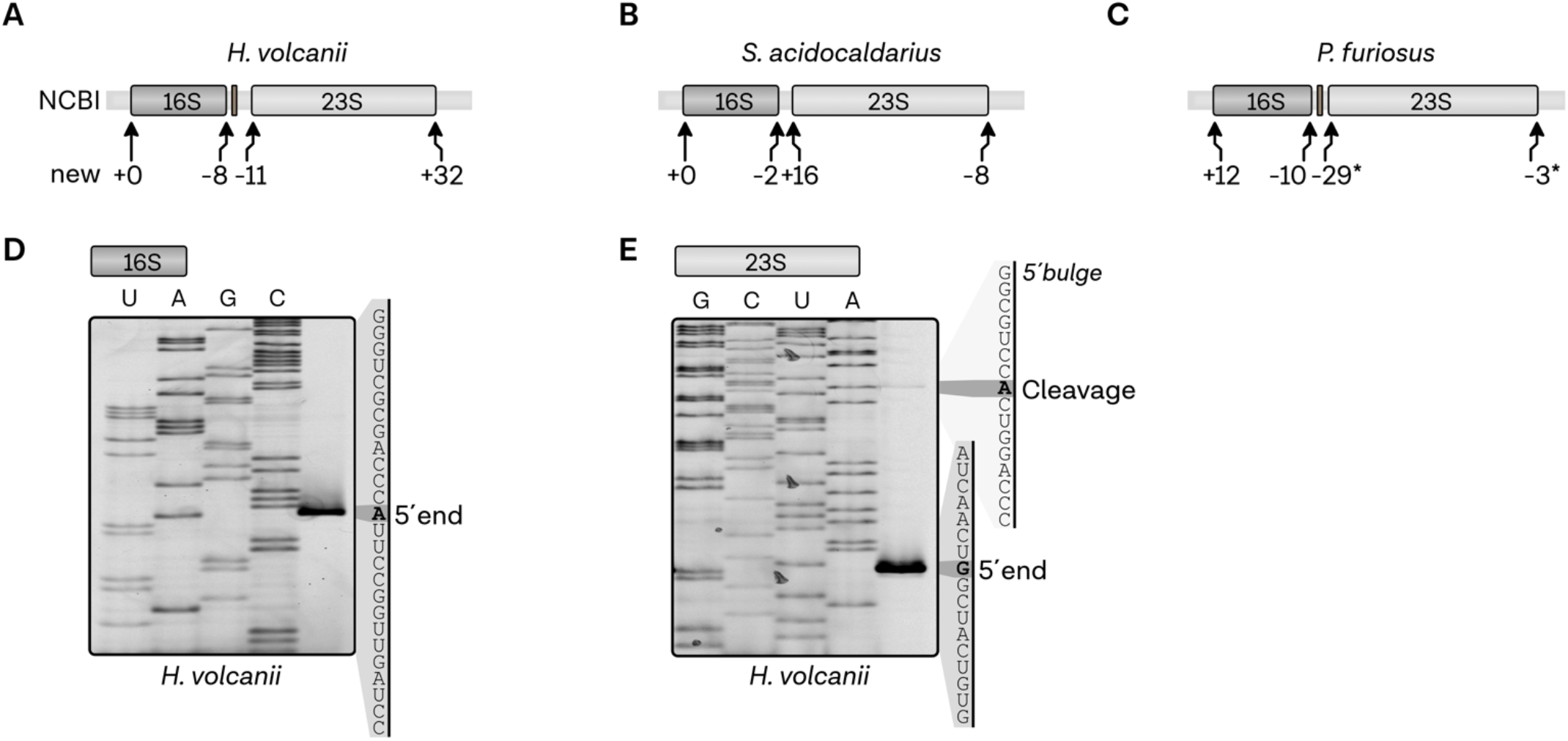
Annotation of mature rRNAs. **A,** Updated annotation of mature rRNA boundaries in *H. volcanii*,**B,***S. acidocaldarius* and **C,***P. furiosus*. Negative numbers indicate that the new position is upstream of formerly annotated positions in the NCBI database, whereas positive ones are downstream. The special case in the 23S rRNA of *Pyrococcus* was marked with an asterisk. **D,** Mapping of mature 16S and E, 23S rRNA 5’ends by primer extension. Comparison to a sequencing ladder allows the assignment of the positions of mature 16S rRNA and 23S rRNA and the 5’bulge of the 23S rRNA.

**Supplementary Figure 6.**
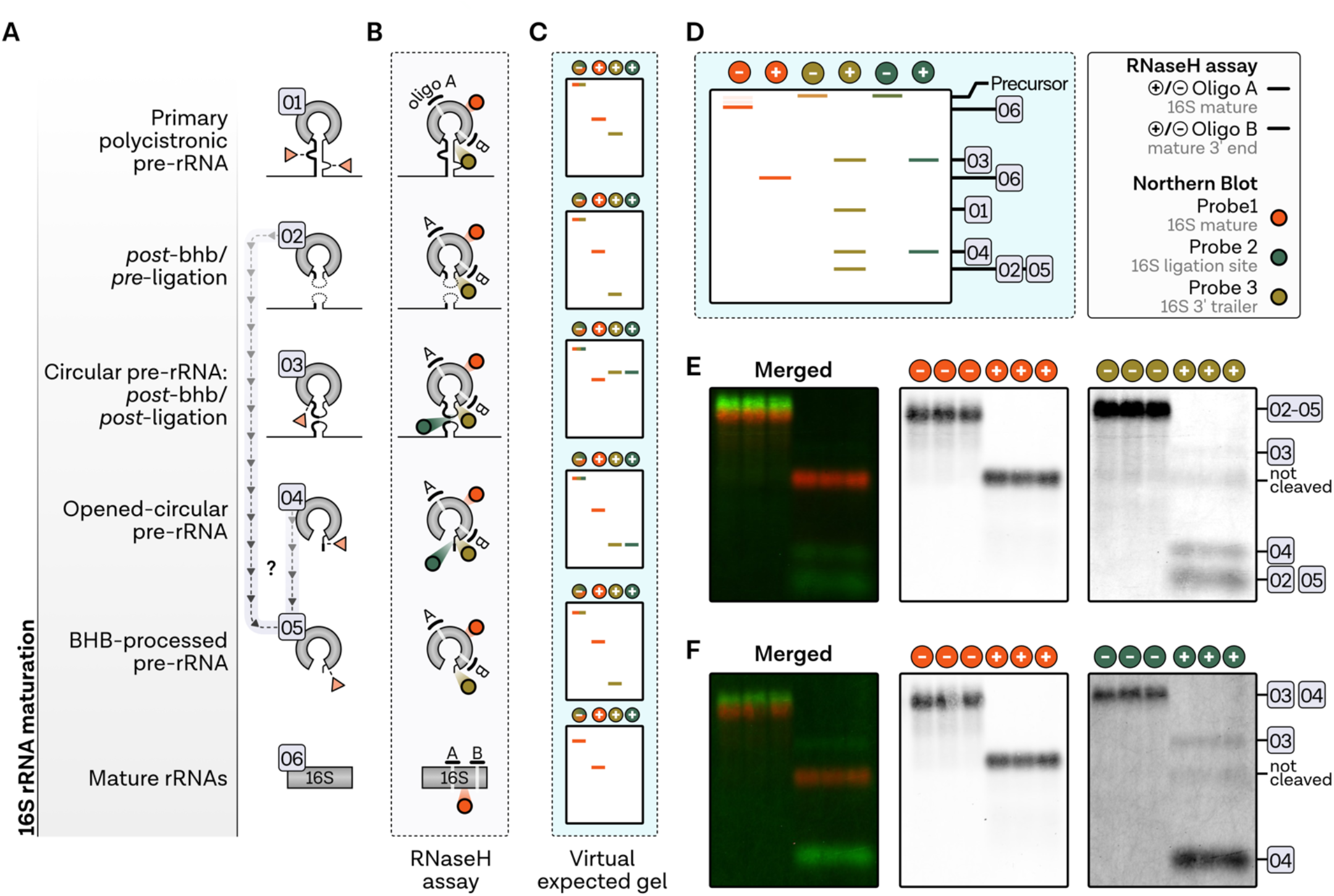
RNaseH cleavage assay. **A,** Proposed 16S rRNA maturation pathway in *Haloferax volcanii*. **B,** Schematic overview of the RNaseH cleavage assay to detect rRNA intermediates. Antisense oligos A and B used for RNAse H-dependent cleavage are shown as black lines, while target regions of the fluorescently labeled Northern Blot probes are highlighted by coloured circles. **C,** Virtual expected gel of each stage (1–6) with (+) and without addition of RNaseH. Probes and expected gel signals are color coded according to the probes shown in panel **D,** which summarises the expected signals using total RNA as input for the RNase H cleavage assay. **E,** Single and merged signals for samples 1 and 3 and F, probes 1 and 2.

**Supplementary Figure 7.**
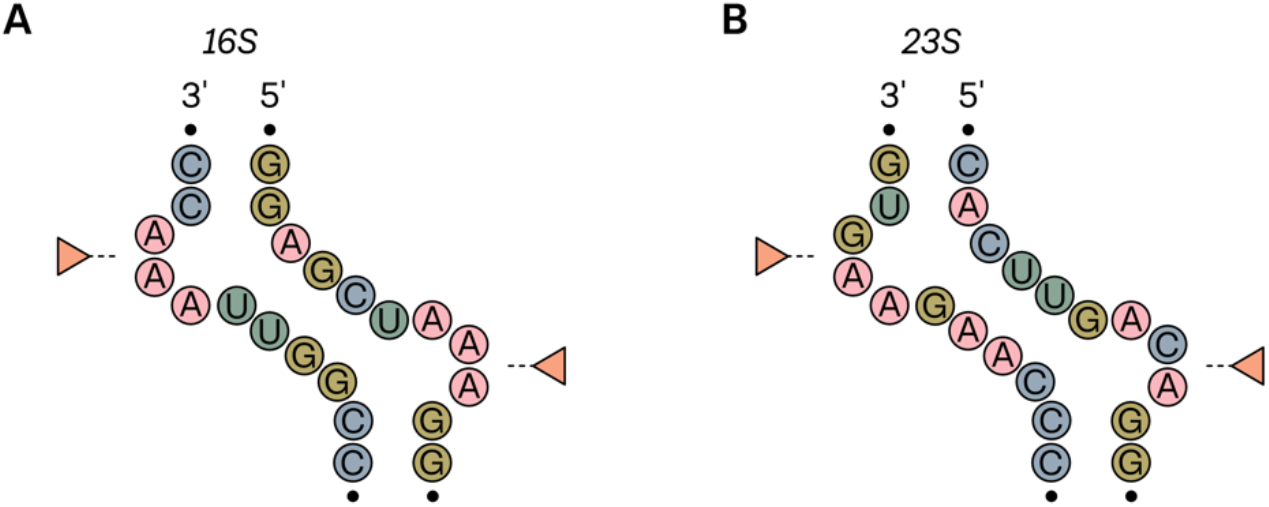
Secondary structure prediction of processing stems in *P. furiosus*. **A,** Secondary structures of the 16S and **B,** 23S rRNA of *P. furiosus*. Cleavage sites are indicated by an orange triangle.

**Supplementary Figure 8.**
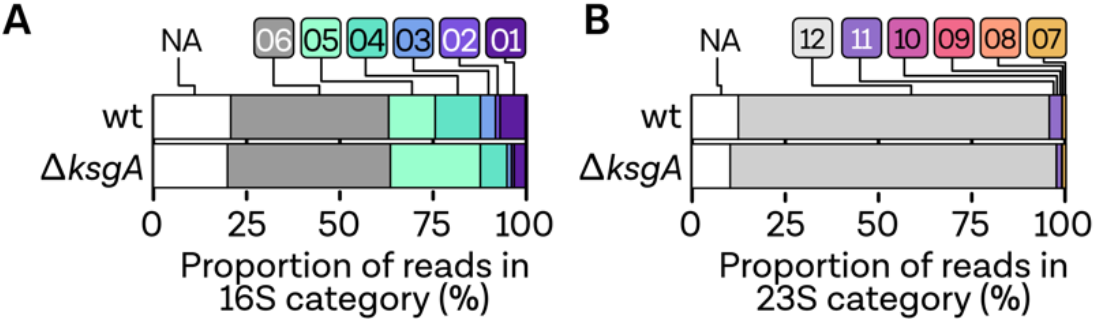
Relative quantification of rRNA intermediate states using direct RNA sequencing data in *Haloferax volcanii*. **A,** The proportion of reads of the different rRNA processing states in the 16S groups and the **B,** 23S read groups are shown as stacked bar charts for the wildtype (wt) and the Δ*ksgA* sample.

